# Uncovering novel pathways for enhancing hyaluronan synthesis in recombinant *Lactococcus lactis*: Genome-scale metabolic modelling and experimental validation

**DOI:** 10.1101/244699

**Authors:** Abinaya Badri, Karthik Raman, Guhan Jayaraman

**Author notes:** Department of Chemical and Biological Engineering, Rensselaer Polytechnic Institute, NY 12180, USA.

## Abstract

Hyaluronan (HA) is a naturally occurring high-value polysaccharide with important medical applications. HA is commercially produced from pathogenic microbial sources. HA-producing recombinant cell factories that are being developed with GRAS organisms are comparatively less productive than the best natural producers. The metabolism of these recombinant systems needs to be more strategically engineered to achieve significant improvement. Here, we use a genome-scale metabolic network model to account for the entire metabolic network of the cell to predict strategies for improving HA production. We here analyze the metabolic network of *Lactococcus lactis* adapted to produce HA, and identify non-conventional overexpression and knock-out strategies to enhance HA flux.

To experimentally validate our predictions, we identify an alternate route for enhancement of HA synthesis, originating from the nucleoside inosine, which has the capacity to function in parallel with the traditionally known route from glucose. Adopting this strategy resulted in a 2.8-fold increase in HA yield. The strategies identified and the experimental results show that the cell is capable of involving a larger subset of metabolic pathways in HA production. Apart from being the first report of the use of a nucleoside to improve HA production, our study shows how experimental results enable model refinement. Overall, we point out that well-constructed genome-scale metabolic models could be very potent tools to derive efficient strategies to improve biosynthesis of important high-value products.

## 1. Introduction

Hyaluronan (HA) is a linear glycosaminoglycan with alternating glucuronic acid and N-acetyl glucosamine repeating units. Due to its very high water-binding capacity and high viscoelasticity, HA assumes multiple roles such as cushioning, lubrication, shock absorbance at important joints and moisture retention in the skin. These make it indispensable for the normal functioning of most organisms. HA is also used in cosmetics as a moisturizer, as a visco-supplement for rheumatoid arthritis patients (Balazs and Denlinger, 1993), in ophthalmic surgeries to replace the lost vitreous fluid (Denlinger and Balazs, 1980), and as a hydrogel for drug delivery systems. A comprehensive review of the uses of HA is discussed elsewhere (Kogan et al., 2006).

Currently, HA is being produced industrially using microbial sources. The general pathway for microbial HA biosynthesis is illustrated in Fig. 1. Native producers like *Streptococci* group A and C that synthesize this polymer as part of their cell capsule have been largely explored for HA production (Chen et al., 2009; Gao et al., 2006; Liu et al., 2009; Shah et al., 2013). More recently, many recombinant platforms have been constructed for HA production, to overcome pathogenicity issues associated with the native producers (Liu et al., 2011). However, it is noteworthy that only one of these recombinant systems (*Bacillus subtilis*) has been scaled up for commercial production (Jia et al., 2013). Widespread application of recombinant cell factories for HA production has been largely impeded by poor yields and low molecular weight of HA, that compares poorly with those from native *Streptococci*.

**Figure 1:**
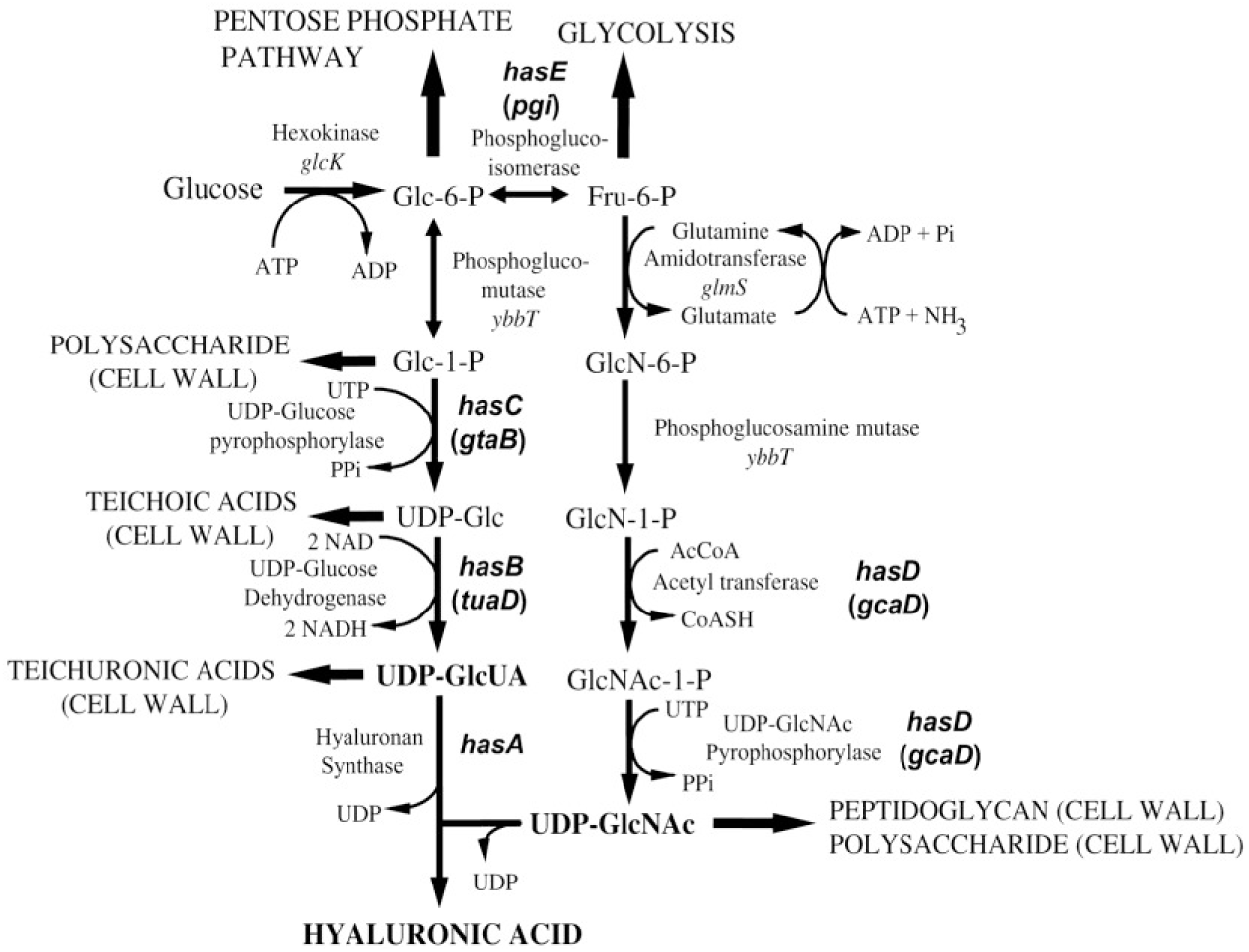
Traditional HA biosynthetic pathway in microorganisms (Widner et al., 2005).

*Lactococcus lactis*, used widely in the production of fermented food products, has a relatively simple metabolism and a long history of safe use. This renders it an ideal chassis for production of medical-grade or food-grade chemicals. HA-producing recombinant *L. lactis* strains, previously developed by our group (Prasad et al., 2010, 2012), have been investigated to identify metabolic factors that affect HA yield (Badle et al., 2014; Chauhan et al., 2014; Kaur and Jayaraman, 2016; Sanghe, 2012). However, only the immediate carbon-backbone based on the traditional HA biosynthetic pathway (Fig. 1) has been considered as the entire canvas for modifications. On the contrary, the metabolic capabilities of the cell can be more effectively understood by a systematic analysis of its entire metabolic network. In this study, we illustrate the potential of genome-scale metabolic network modelling to identify strategies that will guide the metabolic engineering of this organism towards HA production. Recent studies have underlined the importance of such systems-level approaches to re-engineer cellular metabolism (Lee et al., 2012).

Genome-scale metabolic models (GEMs) comprise all the metabolic genes and reactions known to be present in an organism and thus represent the stoichiometry of its metabolic capability. They have been used to rationally identify metabolic engineering strategies while taking the entire network into consideration. A plethora of methods to identify metabolic engineering strategies have also been developed (Burgard et al., 2003; Choi et al., 2010; Hädicke and Klamt, 2010; Kim and Reed, 2012; Pharkya, 2004; Zomorrodi and Maranas, 2012).

For *L. lactis*, three GEMs have been previously curated, and the current study builds on those. Oliveira and co-workers manually reconstructed the first GEM from the annotated genome of *L. lactis* IL1403 (Oliveira et al., 2005). Later, Westerhuis and co-workers reconstructed the *L. lactis* MG1363 model (Verouden et al., 2009) using the semi-automated network reconstruction method, AUTOGRAPH (Notebaart et al., 2006). An updated version of this model, developed by de Vos and co-workers, is the most recent *L. lactis* GEM available (Flahaut et al., 2013). Here we adapt the Flahaut model and analyze it systematically, using a combination of *in silico* approaches, for enhancing HA production. We report several over-expression and knock-out targets. By studying the metabolic sub-network associated with HA and analysis of the strategy that has the maximum effect on HA flux, we identify the supplementation of inosine as a key strategy for increasing HA production in *L. lactis*. We experimentally validate this prediction by demonstrating a 2.8-fold increase in HA production due to the addition of 4 g/L inosine to the *L. lactis* fermentation medium.

## 2. Methods

### 2.1 Computational Methods — Models, Environments and Toolboxes used

The model used in this study was adapted from the *L. lactis* MG1363 GEM published in 2013 (Flahaut et al., 2013). The GEM was analyzed using COBRA toolbox v2.0 (Schellenberger et al., 2011) in the MATLAB R2012b environment (Mathworks Inc., USA).

Simulations involved performing Flux Balance Analysis (FBA) (Varma et al., 1994) or Minimization of metabolic adjustment (MoMA) (Segre et al., 2002) on the entire network. The simulations were subject to steady-state mass balance constraints, experimentally determined bounds on reaction fluxes and additional conditions for knock-outs. The objective function was assigned as biomass flux or HA flux, according to the purpose of the simulation. In cases where HA flux was maximized, a minimum biomass flux was fixed in order to avoid the unrealistic metabolic state of HA production without any cell growth (explained in detail in Supplementary Methods §B.1). Basic operations like adding and removing reactions, changing objective functions, computing steady-state flux distribution using FBA/MoMA were performed using in-built functions in the COBRA toolbox.

Over-expression analysis was done using the concept of Flux Scanning based on Enforced Objective Flux (FSEOF) (Choi et al., 2010). FSEOF identifies a subset of the metabolic network that represents reaction fluxes that increase with an increase in product flux. The FSEOF sub-network visualization was created using Escher {http://escher.github.io/; (King et al., 2015)} and the JSON model required for it was generated using Python 2.7.2 and COBRAPy.

The COBRA toolbox’s built-in function to perform OptKnock (Burgard et al., 2003) was used to identify knock-outs that increase the theoretical maximum product flux. Other knock-outs that move product flux closer to its theoretical maximum were identified by comparing the flux distribution under biomass and product maximization conditions (Supplementary Methods §B.2 and §B.3).

### 2.2 Experimental Methods – Strains, Media, Chemicals and Estimation Protocols

#### Strains and Plasmids

The recombinant HA producing strain used in this study, SJR6, contains the nisin-inducible plasmid pNZ8048 (with chloramphenicol resistance gene) bearing three HA pathway genes viz. Hyaluronan Synthase, UDP glucose-6-dehydrogenase and N-acetyl glucosamine pyrophosphorylase (Prasad et al., 2012).

#### Media components and culture conditions

Inosine was purchased from Sisco Research Laboratories (Mumbai, India). All other media components were procured from HiMedia Laboratories (Mumbai, India). The bioreactor medium consisted of 20 g/L glucose (autoclaved separately), 5 g/L yeast extract, 5 g/L brain heart infusion, 0.5 g/L ascorbic acid, 0.25 g/L MgSO_4_, 1.5 g/L Na_2_HPO_4_, 0.5 g/L KH_2_PO_4_ and 10 µg/mL chloramphenicol. Inosine solutions of the required concentration were autoclaved separately and added to the media at the start of the fermentation. Batch fermentations (30°C, pH 7, 200 rpm, 1 vvm for aerated runs) with a total volume of 1 L were conducted in triplicates in a 2.5 L INFORS HT Benchtop bioreactor (Infors AG, Bottmingen, Switzerland).

#### Fermentation Analysis

Biomass, glucose, inosine, hypoxanthine and HA were estimated at regular intervals. Biomass was estimated from the optical density of the culture (OD_600_), residual glucose measured by GOD-POD assay and HA by the carbazole assay (Bitter and Muir, 1962). Inosine and Hypoxanthine were estimated by a monolithic Luna C-18 Reversed-Phase HPLC column (Phenomenex Inc., Torrance, CA), using linear gradient elution (Farthing et al. 2007). The standard plots for all the metabolites analyzed and the gradient program used for Reverse Phase HPLC is described in detail in Supplementary Methods §A.1.

## 3. Results

We first extended the existing GEM for *L. lactis* to account for the known properties of the strain employed in this study. Subsequently, we performed *in silico* predictions of metabolic engineering strategies to over-produce HA. Based on our *in silico* simulations, we predicted that the supplementation of inosine in the growth medium has a potential to amplify the production of HA. Finally, we validated our predictions through experiments, by demonstrating that there is a 2.8-fold improvement in the production of HA, due to supplementation of inosine in the growth medium. Overall, our results demonstrate the role of systematic model-guided metabolic engineering as a powerful tool for strain improvement.

### 3.1 *In silico* HA production using an adapted *L. lactis* GEM

We adapted the most recent GEM of *L. lactis* (Flahaut et al., 2013) to investigate the metabolic capabilities of the organism towards HA production. The model in SBML format is available as Supplementary File S1. First, we added the reaction catalyzed by HA synthase to match the metabolism of the recombinant strain. Next, we replaced the flux bounds of uptake reactions specified in the original model with those that have been reported for the strain we used (Badle et al., 2014). We eliminated gaps and blocked reactions from the model, which resulted in a version of the model specific for the nutrient sources supplied in this study. We also included an additional constraint that enabled homolactic fermentation (Gao et al., 2006). A detailed description of the changes made to the model is available in Supplementary Methods §A.2.

### 3.2 New metabolic engineering strategies for HA production in *L. lactis*

We analyzed the GEM described above to identify strategies commonly employed to engineer the metabolism of an organism towards a product of interest. The approaches we employed can be grouped into two categories: (1) amplifying fluxes of reactions that aid in HA production; (2) knocking-out or reducing the flux of reactions that compete with HA. Model analysis in the first approach revealed very interesting results, one of which experimentally improved HA production by 2.8 fold. This will be discussed in detail in the rest of the article. On the other hand, the latter approach led to intuitive/previously reported strategies that further establish the validity of our methodology. They have been discussed in detail in Supplementary Results §B.2 and §B.3

#### 3.2.1 Over-expression Strategies

Over-expression of genes in the product pathway has been employed in several cases of metabolite production (Farmer and Liao, 2000; Shimada et al., 1998). We use the concept of FSEOF (Choi et al., 2010) to systematically obtain an exhaustive set of all candidate targets for over-expression from the metabolic network of *L. lactis* (Table S1). FSEOF identifies those reactions, which tend to carry higher flux whenever product flux increases (as mandated by constraints in the simulations), as observed by simulating the metabolic network using FBA. These reactions are then evaluated as potential candidates for overexpression.

We thus identified 78 reactions (40 reversible and 38 irreversible reactions) that included the traditional HA pathway reactions. The involvement of glutamine in HA pathway could be a possible reason for the presence of some amino acid biosynthesis reactions on the list. Apart from these, several nucleotide metabolism reactions came up on the list. They appeared to contribute to the carbon backbone of HA. Fig. S1 shows a visualization of the reaction sub-network constructed out of this list of reactions, using Escher (King et al., 2015). This schematic helps to illustrate how these reactions contribute to HA.

Though all the reactions in the sub-network are indeed contributing to increasing HA in some way or another, it would be ideal to point out those which would be better targets to guide metabolic engineering. From the FSEOF analysis, this study brings out two types of over-expression targets (Fig. 2). The first type of over-expression targets are the obvious pathway enzymes. This list consists of eight reactions of the HA pathway *viz.* Hyaluronan synthase, UDP-glucose dehydrogenase, UDP-N-acetylglucosamine diphosphorylase, UTP glucose-1-phosphate uridylyltransferase, Glucosamine-1-phosphate N-acetyltransferase, phosphoglucosamine mutase, phosphoglucomutase and Glutamine fructose-6-phosphate transaminase. Many of these have already been verified experimentally in previous studies in *L. lactis* (Prasad et al., 2010, 2012; Zhang et al., 2016).

**Figure 2:**
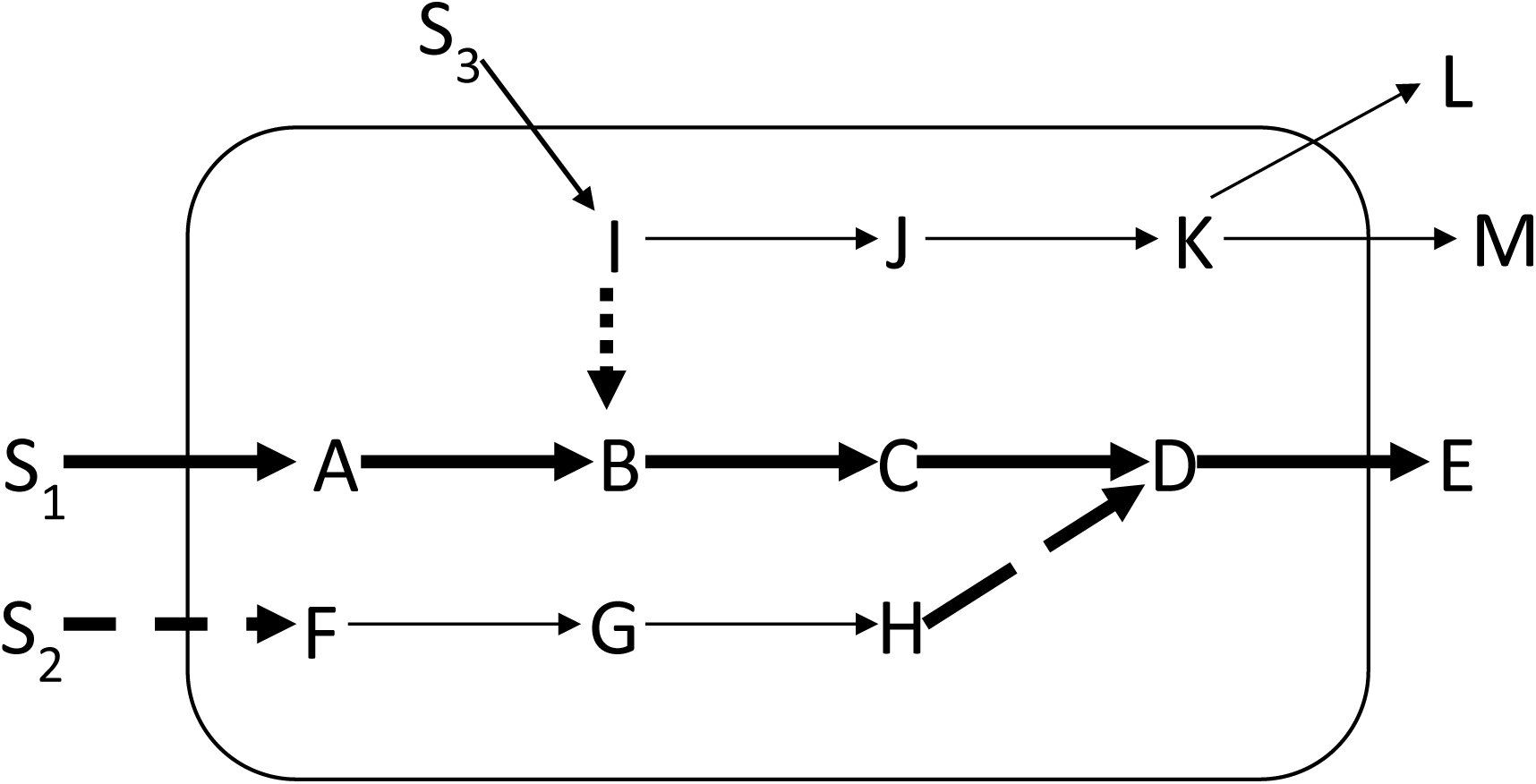
Categories of over-expression targets in the product sub-network of the cell: Type I targets (bold solid lines) from the traditional biosynthetic pathway; Type II-A targets (dotted lines) involve branch nodes and can be identified directly. Type II-B targets (dashed lines) can be traced out from contributors (long dashed lines) that are parts of a linear path.

The second type of over-expression targets predicted herein comprises *non-intuitive* targets. These have been further classified, based on whether they are part of a branch or a linear pathway. The first category includes those reactions whose substrates are branch nodes. When these are over-expressed, the fraction of the branch point metabolite going towards the pathway of interest will increase, making them good candidates for over-expression. Such reactions in the FSEOF list were identified by searching for important branch nodes (non-currency metabolites participating in more than two active reactions) in the FSEOF sub-network. This analysis pointed to five enzymes as targets for over-expression (Table 1).

**Table 1:** Effect of each predicted Type-II over-expression strategy on HA flux

| Strategy | Slope of maximum HA vs. substrate<br>(mol HA/mol substrate of strategy) |
| --- | --- |
| **Type-II(A)** |  |
| 2-deoxyribose-5-phosphate aldolase | 6.234 |
| Glutamine synthase | 1 |
| Acetaldehyde dehydrogenase (acetylating) | 0.659 |
| L-serine deaminase | 0.564 |
| Deoxycytidine deaminase | 0.25 |
| **Type-II(B)** |  |
| Glutamine uptake | 1 |
| Inosine uptake | 0.25 |
| Serine uptake | 0.065 |
| Isoleucine uptake | 0.03 |
| Aspartate uptake | 0.027 |

The second category of type-II targets stems from reactions that are part of linear pathways in the sub-network. Over-expression of these enzymes might not increase product flux since reactions preceding it and succeeding it would still have unchanged fluxes. Here, we reasoned out that first reactions of the linear path would be better targets than an intermediate reaction. However, the starting points to a linear bit in the pathway could lead to either branch nodes or exchanges (I or S_2_ in Fig. 2). The case of branch point origins has been handled in the first category of type II targets. To identify exchange start points, the exchange reactions involved in the HA sub-network were examined. There were totally 13 exchange fluxes in the FSEOF list. The 5 uptakes out of these have been put forth as over-expression targets of this class (Table 1). One of these targets, inosine, has been experimentally validated in this study and is discussed in detail in the following section.

### 3.3 Experimental Validation of Predicted Strategy

A total of 18 overexpression and 12 knock-out targets have been put forth by the model analyses conducted in this study. This list of strategies is not exhaustive but covers major types of strategies to engineer the metabolism towards HA production. Some of these strategies have been experimentally observed to be effective in previous studies. For example, the traditional *has* operon genes have been cloned in various combinations in *L. lactis* to increase HA production (Chauhan et al., 2014; Chien and Lee, 2007; Prasad et al., 2010). Apart from the operon, other pathway genes like phosphoglucomutase have been over-expressed (Zhang et al., 2016). Knocking out lactate dehydrogenase gene has already been shown to bring about increase in HA yield and molecular weight (Kaur and Jayaraman, 2016). Among the type-II targets for over-expression put forth in this study, though none have been actually tested in practice, there are some reports on the increase in HA yield upon amino acid addition in *Streptococci* (Shah et al., 2013).

Importantly, this study suggests, for the first time, the use of nucleosides to increase HA flux. Inosine, a nucleoside containing hypoxanthine (nitrogen base) and ribose, was one among the five metabolites whose uptake rate was found to increase in the model with a forced increase in HA production rate (Table 1). This indicates that increasing the inosine uptake rate in the cell might lead to an increase in HA flux. Apart from this, out of the 10 new over-expression targets in Table 1, the one with the highest increase in theoretical HA flux was the 2-deoxyribose-5-phosphate aldolase reaction. This is one of the downstream reactions of inosine catabolism according to the sub-network in Fig. S1, suggesting that increasing the uptake of inosine would be one of the most effective strategies put forth in this study. Hence, this strategy presents itself as an interesting one among the others and was therefore tested for improved HA production.

#### 3.3.1 Addition of Inosine resulted in Hypoxanthine secretion in recombinant *L. lactis*

Hypoxanthine was observed to accumulate in the culture media that were supplemented with inosine. This was identified in the chromatogram used to estimate the residual inosine concentration (Fig. S2). The concentration profiles of hypoxanthine accumulated and inosine consumed (Fig. 3) show that amounts of hypoxanthine accumulated in the medium were equimolar to the consumption of inosine. The equimolar secretion of hypoxanthine indicates that the inosine taken up from the medium is cleaved and the nucleobase is secreted back into the medium. This has been reported in certain other organisms too (Bzowska et al., 2000; He et al., 1994). Hence, what remains in the cell is the ribose moiety from the nucleoside.

**Figure 3:**
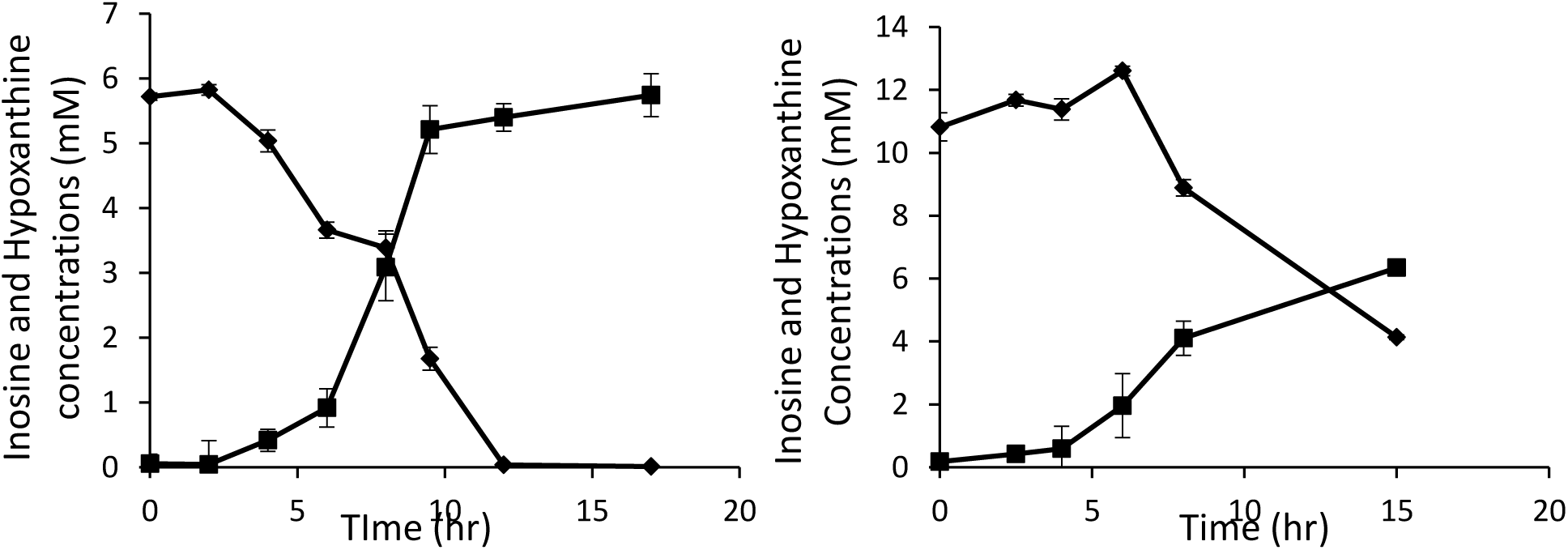
Equimolar Hypoxanthine (■) production with inosine (♦) consumption (a) for 1.5 9 g/L inosine and (b) 3 g/L inosine runs

#### 3.3.2 Inosine uptake led to a 2.8-fold increase in HA yield from total substrate supplied

A positive correlation was observed between the initial concentration of inosine added to the medium and the final HA concentration obtained in the fermentation (Fig. 4). Compared to the control run without inosine addition (0.37 g/L HA), the final HA concentration obtained increased 3-fold (to ∼1.1 g/L) when 4g/L inosine was added to the medium. The HA yield per carbon-mole (C-mole) of the substrate (glucose + inosine) also increased 2.8-fold for the batches supplemented with inosine as shown in Table 2.

**Table 2:** Yield of HA from inosine and glucose in the batch bioreactor runs.

| Parameter | Control | Initial Inosine<br>1.5 g/L | Initial Inosine<br>3 g/L | Initial Inosine<br>4 g/L |
| --- | --- | --- | --- | --- |
| ΔGlucose (g/L) | 20 | 20 | 20 | 20 |
| C-moles of Glucose | 0.667 | 0.667 | 0.667 | 0.667 |
| Average ΔInosine (g/L) | 0 | 1.5 | 2 | 2.5 |
| C-moles of inosine | 0 | 0.056 | 0.075 | 0.093 |
| Average HA titer (g/L) | 0.37 | 0.53 | 0.64 | 1.1 |
| C-moles of HA monomer | 0.013 | 0.019 | 0.023 | 0.041 |
| $Y_{HA/S\_total}$ (C-mole/C-mole) | 0.019 | 0.0262 | 0.031 | 0.054 |

**Figure 4:**
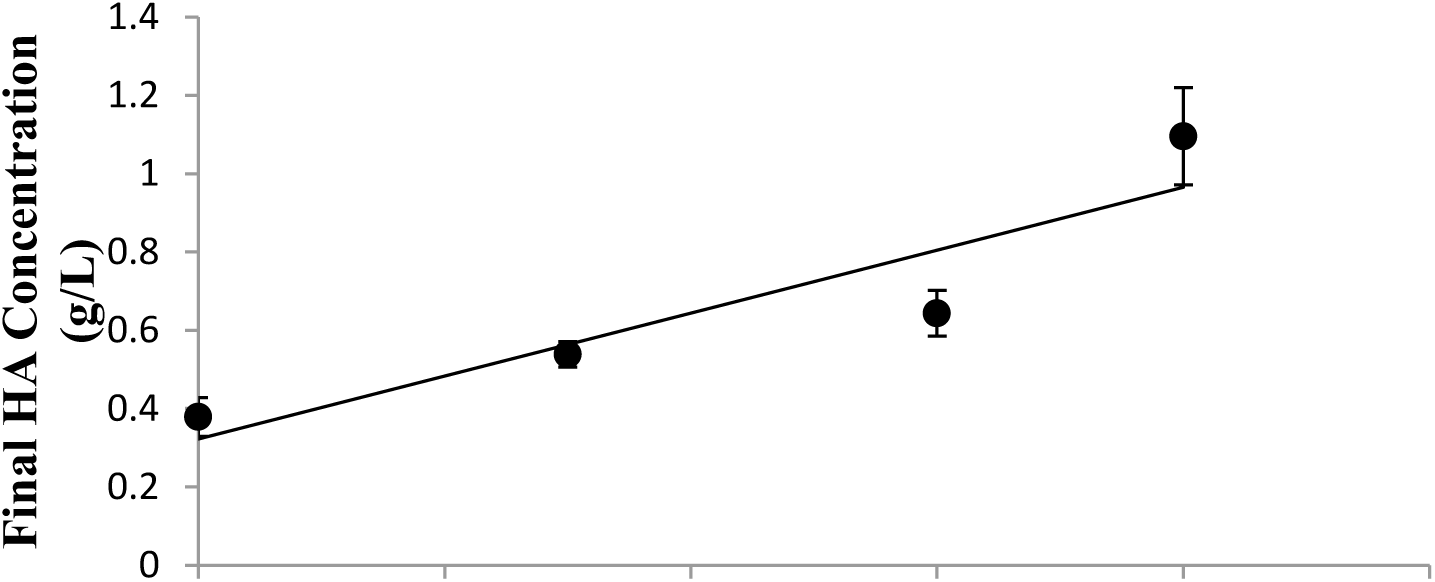
Batch Experiments in Bioreactors. Final HA concentration vs. initial inosine concentration in batch bioreactor study with 20g/L glucose

#### 3.3.3 Higher initial glucose concentration helped overcome the incomplete consumption of inosine

The fermentation profiles for the batch bioreactor runs indicate that inosine utilization is incomplete in the runs with higher initial inosine concentration, unlike the runs with 1.5 g/L inosine (Fig. 5). Inosine consumption appeared to stop after complete consumption of glucose (onset of stationary phase). Considering that the inosine consumption aids HA production, it would be beneficial to have complete consumption of inosine in the batch fermentation.

**Figure 5:**
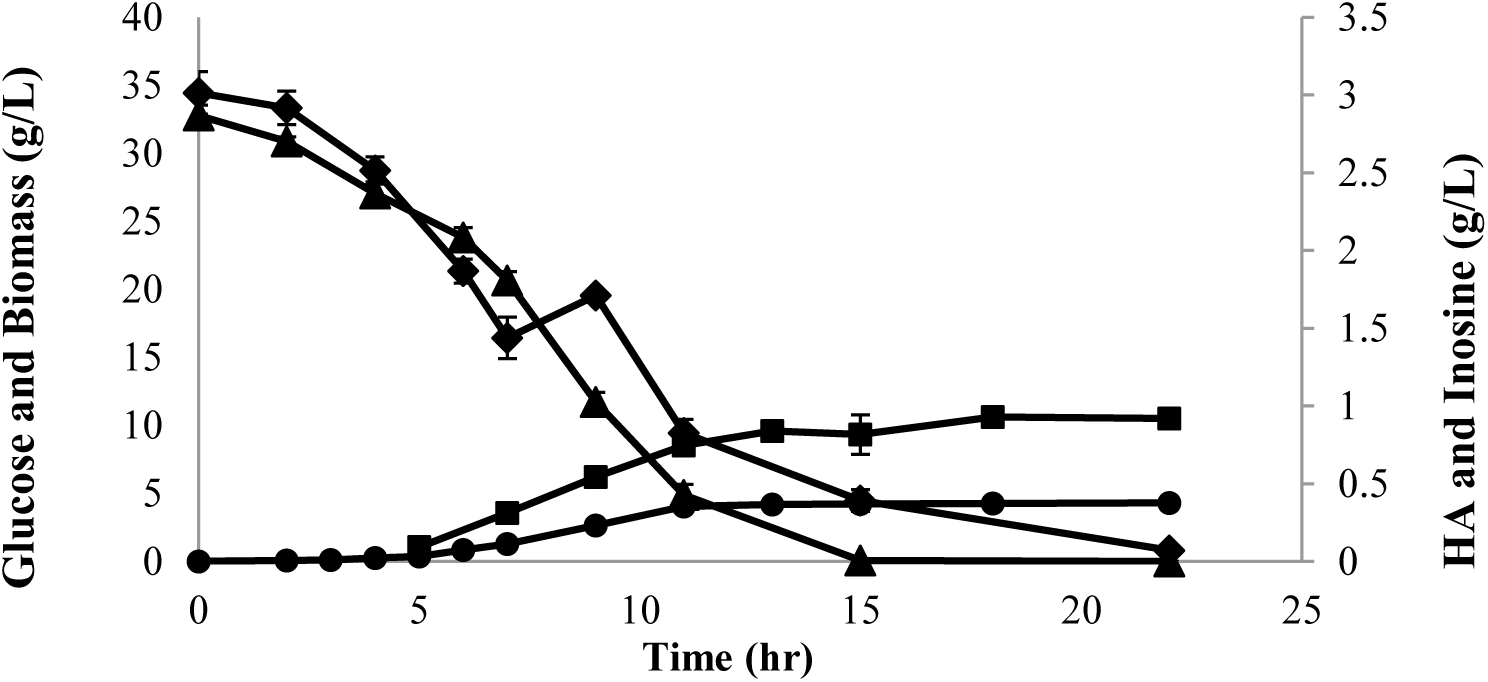
Fermentation profile of batch run with 3 g/L inosine and 30 g/L glucose showing higher inosine consumption (glucose (▲); biomass (●); HA (■); inosine (♦))

To enhance inosine consumption, initial glucose levels were increased to delay glucose exhaustion (and hence delay the onset of stationary phase). Higher initial glucose level (30 g/L) for the same level of inosine enabled complete consumption of inosine (Fig. 6). A correspondingly high concentration of HA (0.97 g/L) was obtained from this fermentation process.

**Figure 6:**
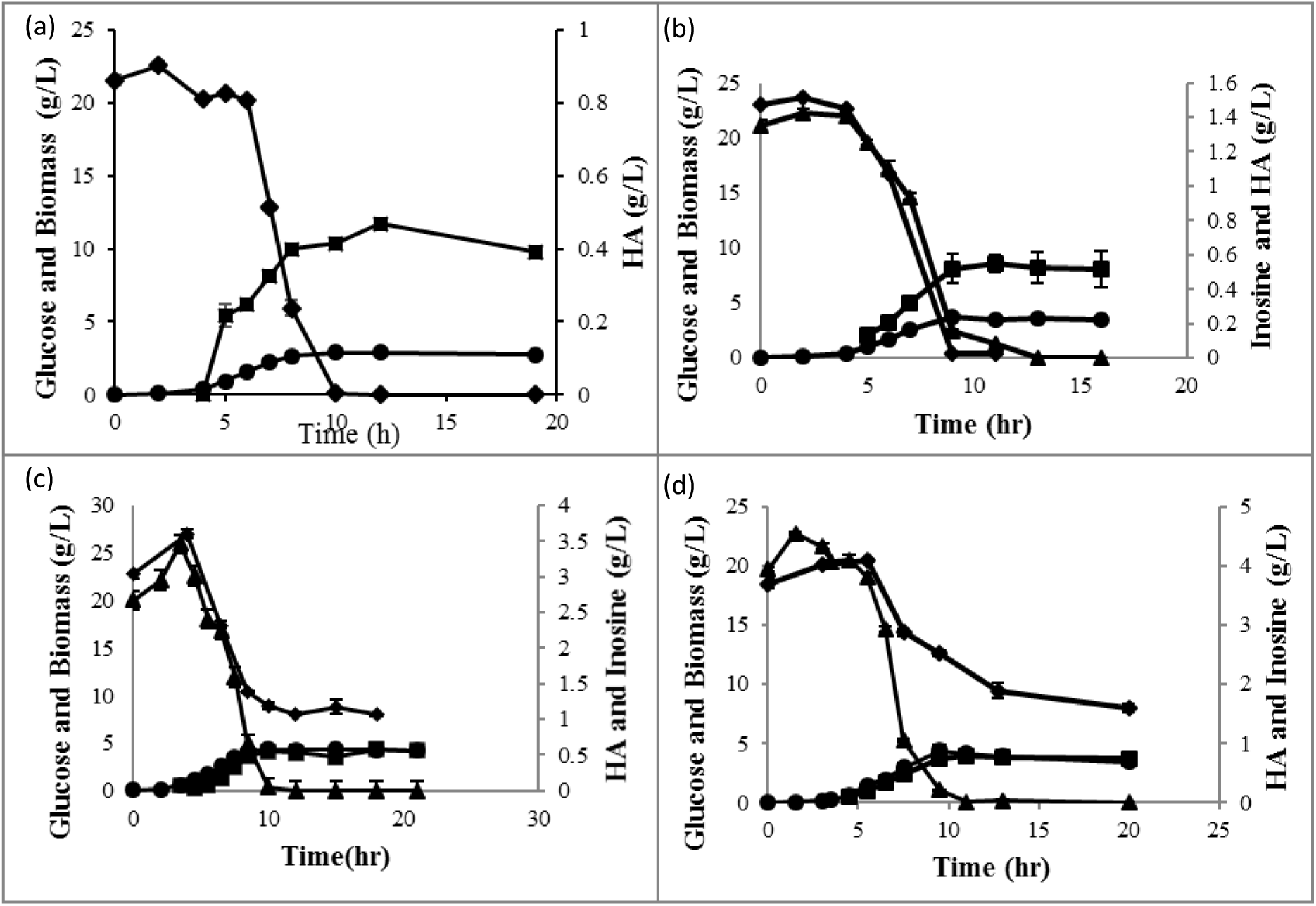
Fermentation profiles of batch run with 20 g/L glucose and 0 g/L (a), 1.5 g/L (b), 3 g/L (c) and 4 g/L (d) inosine. Unconsumed inosine observed in (c) and (d). glucose (▲); biomass (●); HA (■); inosine (♦)

### 3.4 Extrapolation of The Predicted Strategy

Setting maximization of HA flux as the FBA objective ideally picks the best route to synthesize HA from the network. This (represented in Fig. S1) shows that inosine is transported into the cell and further phosphorylated to cleave the ribose sugar off the nucleoside by purine nucleoside phosphorylase. The hypoxanthine nucleobase generated is further metabolized by Hypoxanthine Phosphoribosyl Transferase (HXPRT) and adenylosuccinate lyase. The ribose-1-phosphate generated in this process is transferred to uracil, forming uridine, which then goes through a cycle of kinases and phosphatases and is also reduced to give the corresponding deoxy-ribose sugar. The deoxy-ribose generated is finally cleaved by 2-deoxyribose phosphate aldolase that produces acetaldehyde and glyceraldehyde-3-phosphate. Both these compounds aid the HA pathway via acetyl-CoA generation and gluconeogenesis respectively.

#### 3.4.1 Identification of Hypoxanthine Exporter Gap Enabled Model Improvement

Hypoxanthine secretion into the medium indicates that *L. lactis* did not metabolize the hypoxanthine further contrary to model prediction. Though nucleobase secretion has been observed in several other reports available in literature, the FSEOF analysis predicted re-routing of hypoxanthine to inosine monophosphate by HXPRT instead. This reaction requires PRPP. Furthermore, the inosine monophosphate is converted to cyclic-AMP by adenylosuccinate lyase. This reaction requires both L-alanine and GTP. The cyclic-AMP breaks into AMP and fumarate, that is ultimately converted to succinate and secreted out.

This path is more expensive compared to just secreting hypoxanthine as in the real system. Upon probing the model to find out why such a route was predicted, we identified that this was due to the absence of a hypoxanthine exporter in the model. When this transport reaction was added, hypoxanthine was fluxed out instead of going to HXPRT. The model chose a more complicated path here due to this gap. This knowledge gained from the real system about hypoxanthine secretion was employed in model refinement. Hypoxanthine secretion implies that the inosine strategy in effect works *via* ribose, as mentioned in §3.3.1

#### 3.4.2 A Transaldolase Knock-In Improves the Inosine Strategy

Once in the cell, there are many fates of catabolism to the ribose moiety. According to Ipata and co-workers, the sinks for ribose phosphates in the cell are nucleoside interconversion reactions and pathways that catabolize ribose as a carbon/energy source (Tozzi et al., 2006). The most common way to use ribose as a carbon/energy source is to join the pentose phosphate pathway. Contrarily, converting the ribose to deoxyribose to metabolize it, as shown in the model analysis, seems to be a convoluted way to contribute to HA production.

However, the pentose phosphate pathway (PPP) is incomplete in *L. lactis* due to the absence of the transaldolase enzyme. This could be the reason why the model predicted a path from inosine to HA *via* 2-deoxyribose-5-phosphate aldolase instead of PPP. Calculating a stoichiometric HA yield from this ribose moiety for each of the conditions (Table 3) indicates that fluxing the ribose through the complete PPP leads to the most efficient conversion. Accordingly, a knock-in of transaldolase would help in the full realization of the inosine addition strategy by facilitating the synthesis of one HA monomer from 2.4 ribose molecules. This was also tested using the model, the results of which are shown in Fig. 7. This way, the model also served as a source for verifying an improvisation to the inosine addition strategy.

**Table 3:**
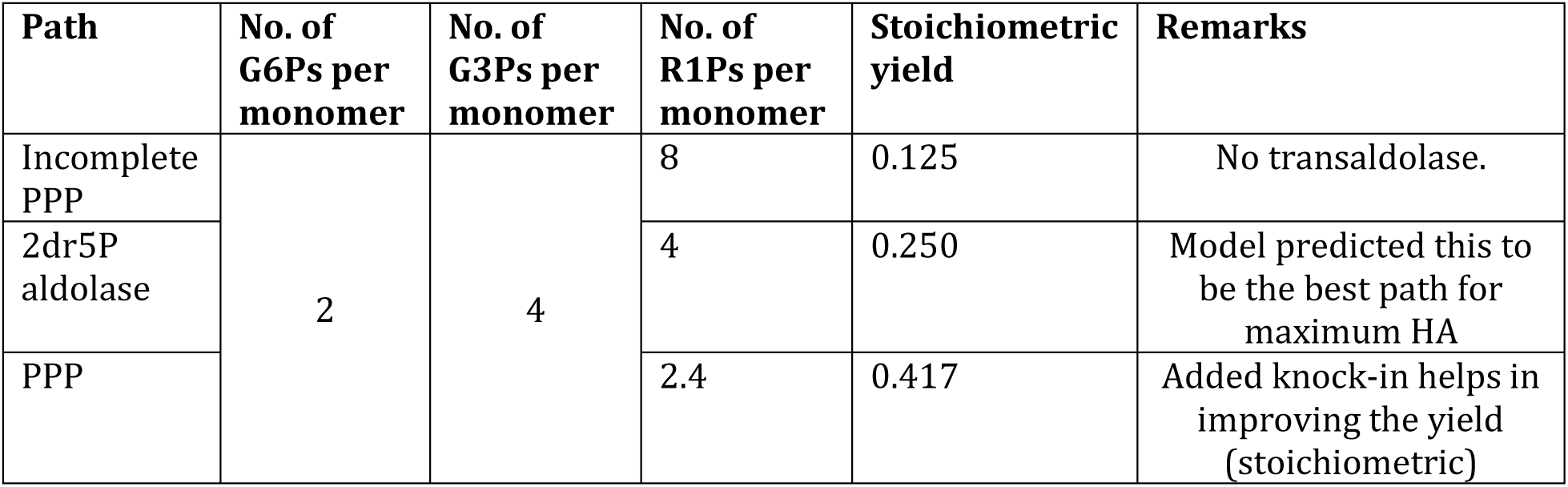
Stoichiometric yield of HA from ribose phosphate of inosine through different paths. (PPPpentose phosphate pathway; G6Pglucose-6-phosphate; G3Pglyceraldehyde-3-phosphate; R1Pribose-1-phosphate)

| Path | No. of G6Ps per monomer | No. of G3Ps per monomer | No. of R1Ps per monomer | Stoichiometric yield | Remarks |
| --- | --- | --- | --- | --- | --- |
| Incomplete PPP | 2 | 4 | 8 | 0.125 | No transaldolase. |
| 2dr5P aldolase |  |  | 4 | 0.250 | Model predicted this to be the best path for maximum HA |
| PPP |  |  | 2.4 | 0.417 | Added knock-in helps in improving the yield (stoichiometric) |

**Figure 7:**
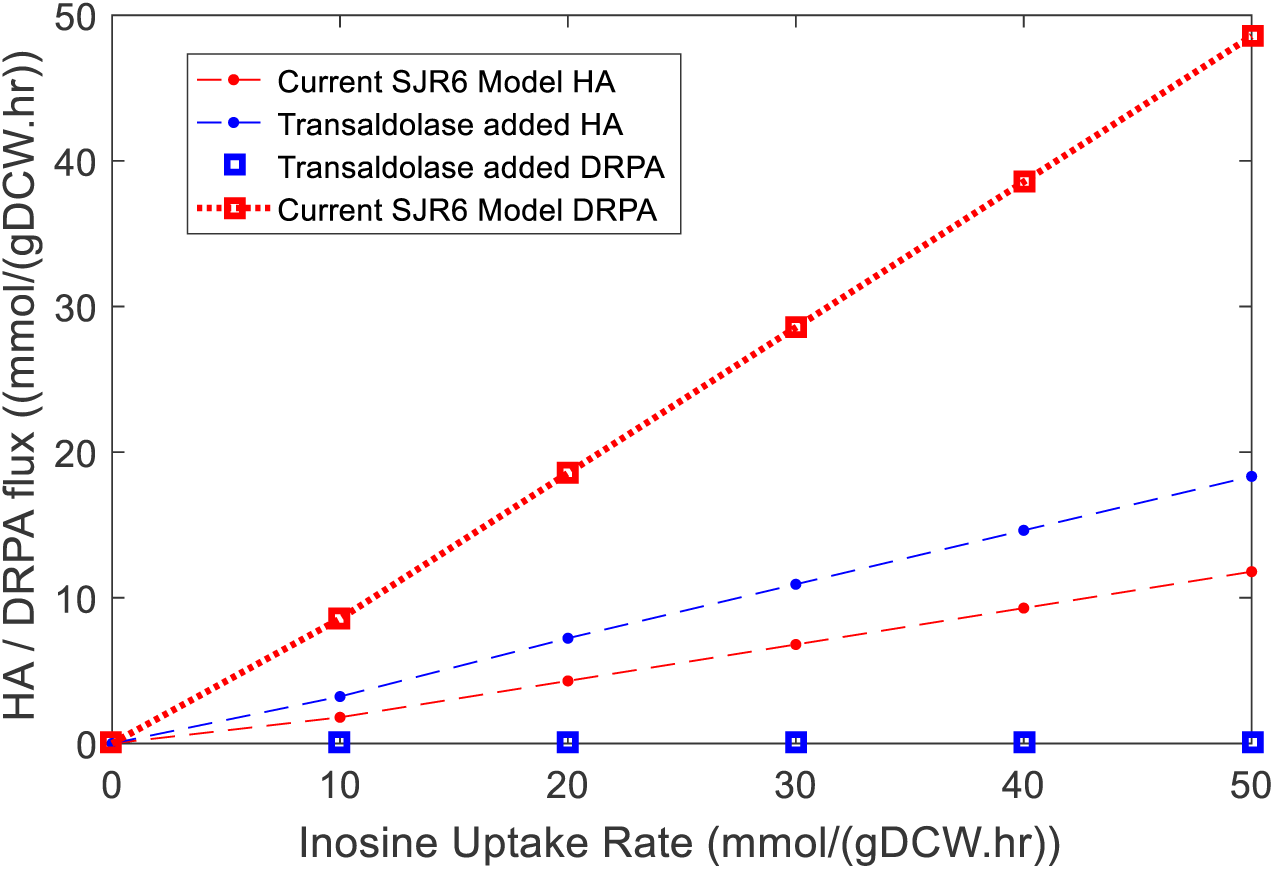
Increase in the slope of HA flux from inosine upon knock-in of transaldolase reaction. (2-deoxyribose-5-phosphate aldolase (DRPA) reaction inactive when transaldolase is present)

## 4. Discussion

Literature reports on HA production have mostly focused on improving process strategies and estimating the sensitivity of the system to external factors (Du, 2009; Duan et al., 2008; Vázquez et al., 2010). A few other studies discuss genetic engineering strategies that confer HA producing capabilities to GRAS organisms that originally did not produce HA (Chien and Lee, 2007; Jeong et al., 2014; Mao and Chen, 2007; Widner et al., 2005). Flux analysis studies have predominantly looked at agreeing with conditions observed in the real system, though some of them have furthered to identify bottlenecks in HA production (Badle et al., 2014; Jagannath and Ramachandran, 2010; Shah et al., 2013). However, these studies have not attempted a systematic identification of metabolic engineering strategies using a genome-scale model of the microorganism. The analysis of the GEM in this study yielded several unexplored and interesting strategies for improving the biosynthesis of HA in *L. lactis*.

This study used FSEOF to delineate the entire sub-network related to product synthesis. Further screening to identify overexpression targets was done by categorizing the locations of the enzymes in the sub-network and then looking at relative variability rather than a flux variability analysis as reported in the original paper by Lee and co-workers (Choi et al., 2010). This study also uncovers an extensive range of new leads from the sample experimental validation section as described in §3.4. Thus, we show that the strategies predicted by these models can be further analyzed thoroughly to get more insights for improvisation.

The updated version of the model, as well as the experiments in this study, suggest that the key contributor to the HA pathway in the inosine strategy is ribose. However, since *L. lactis* cannot catabolize ribose, inosine seems to be an alternate way to get ribose in the cell. In ribose catabolizing organisms, the uptake would be severely limited by the presence of glucose (catabolite repression), leading to a diauxic growth pattern. However, an equivalent strategy in such organisms would be to engineer co-utilization of ribose and glucose. Several strategies to do this, including phosphotransferase transport system deletion (Nichols et al., 2001) and alteration of the regulatory proteins (Yao and Shimizu, 2013) have been employed in model organisms like *E. coli*.

## 5. Conclusions

In this era where GEMs are becoming increasingly popular as tools for model-driven design of experiments (McCloskey et al., 2014; Mienda, 2016), this study exploits the power of analyzing such models for enhancing production of HA. This study is hence the first to systematically identify strategies for HA production from the *L. lactis* metabolic network. It is also the first report, to the best of our knowledge, that considers the entire *L. lactis* network to design strategies for enhanced HA production. These results would guide a more nuanced metabolic engineering of *L. lactis* to overproduce HA.

Contrary to previous focus on the traditional HA biosynthetic pathway, the results reported here indicate that the cell is capable of involving a much a bigger subset of metabolism for producing HA. Apart from identifying a total of 10 new over-expression and nine knock-out targets, this study also experimentally demonstrates a novel route for in HA synthesis. This work, for the first time, reported the use of a nucleoside to enhance HA production. A three-fold increase in HA titer was also observed in batch fermentations supplemented with 4 g/L inosine. We also show that enhancements of the predicted strategies are possible by further model analysis. Based on these results, we postulate that the method used in this study can be applied to predict strategies and improve production of other valuable high-value metabolites from such model biological systems.

