## Supplementary Materials for "Uncovering novel pathways for enhancing hyaluronan synthesis in recombinant *Lactococcus lactis*: Genome-scale metabolic modelling and experimental validation"

### Supplementary Figures and Tables:

**Table S1: 78 Reactions identified by FSEOF**

| S.No. | Reaction Name | Reaction Formula |
|---|---|---|
| 1 | '2 methylbutanal dehydrogenase acid forming ' | 'h2o[c] + nad[c] + 2mbal[c] <=> 2 h[c] + nadh[c] + 2mba[c] ' |
| 2 | '2 methylbutanoic acid transport H symport ' | 'h[c] + 2mba[c] <=> h[e] + 2mba[e] ' |
| 3 | '3 methyl 2 oxopentanoate decarboxylase' | 'h[c] + 3mop[c] -> 2mbal[c] + co2[c] ' |
| 4 | 'acetaldehyde dehydrogenase acetylating ' | 'coa[c] + nad[c] + acald[c] <=> h[c] + nadh[c] + accoa[c] ' |
| 5 | 'acetaldehyde reversible transport' | 'acald[e] <=> acald[c] ' |
| 6 | 'deoxycytidine kinase' | 'atp[c] + dcyt[c] <=> adp[c] + h[c] + dcmp[c] ' |
| 7 | 'adenine transport via proton symport reversible ' | 'h[e] + ade[e] <=> h[c] + ade[c] ' |
| 8 | 'adenylate kinase' | 'atp[c] + amp[c] <=> 2 adp[c] ' |
| 9 | 'adenylosuccinate lyase' | 'dcamp[c] -> amp[c] + fum[c] ' |
| 10 | 'adenylosuccinate synthetase' | 'asp_L[c] + gtp[c] + imp[c] -> 2 h[c] + pi[c] + dcamp[c] + gdp[c] ' |
| 11 | 'adenosylhomocysteine nucleosidase' | 'h2o[c] + ahcys[c] -> ade[c] + rhcys[c] ' |
| 12 | 'alanine racemase' | 'ala_L[c] <=> ala_D[c] ' |
| 13 | 'L alanine transaminase' | 'akg[c] + ala_L[c] <=> pyr[c] + glu_L[c] ' |
| 14 | 'D alanine transport inout via proton symport' | 'h[e] + ala_D[e] <=> h[c] + ala_D[c] ' |
| 15 | 'L aspartate transport in via proton symport' | 'h[e] + asp_L[e] -> h[c] + asp_L[c] ' |
| 16 | 'ATP synthase three protons for one ATP ' | 'adp[c] + pi[c] + 3 h[e] <=> atp[c] + h2o[c] + 2 h[c] ' |
| 17 | 'CO2 transport out via diffusion' | 'co2[e] <=> co2[c] ' |
| 18 | 'CTP synthase NH3 ' | 'atp[c] + nh4[c] + utp[c] -> adp[c] + 2 h[c] + pi[c] + ctp[c] ' |
| 19 | 'cytidylate kinase dCMP ' | 'atp[c] + dcmp[c] <=> adp[c] + dcdp[c] ' |
| 20 | 'deoxycytidine deaminase' | 'h2o[c] + h[c] + dcyt[c] -> nh4[c] + duri[c] ' |
| 21 | 'deoxyribose phosphate aldolase' | '2dr5p[c] -> acald[c] + g3p[c] ' |
| 22 | 'purine nucleoside phosphatase deoxyuridine ' | 'pi[c] + duri[c] <=> ura[c] + 2dr1p[c] ' |
| 23 | '2 methyl butanoic acid exchange' | '2mba[e] <=> ' |
| 24 | '4 Aminobutanoate exchange' | '4abut[e] <=> ' |
| 25 | 'Acetaldehyde exchange' | 'acald[e] <=> ' |
| 26 | 'Adenine exchange' | 'ade[e] <=> ' |
| 27 | 'D Alanine exchange' | 'ala_D[e] <=> ' |
| 28 | 'L Aspartate exchange' | 'asp_L[e] <=> ' |
| 29 | 'CO2 exchange' | 'co2[e] <=> ' |
| 30 | 'L Glutamate exchange' | 'glu_L[e] <=> ' |
| 31 | 'L Isoleucine exchange' | 'ile_L[e] <=> ' |
| 32 | 'Inosine exchange' | 'ins[e] <=> ' |
| 33 | 'L Lactate exchange' | 'lac_L[e] <=> ' |

| | | |
|---|---|---|
| 34 | 'Exchange for Serine' | 'ser_L[e] <=> ' |
| 35 | 'Succinate exchange' | 'succ[e] <=> ' |
| 36 | 'fructose bisphosphate aldolase' | 'fdp[c] <=> dhap[c] + g3p[c] ' |
| 37 | 'fructose bisphosphatase' | 'h2o[c] + fdp[c] -> pi[c] + f6p[c] ' |
| 38 | 'fumarate reductase NADH ' | 'h[c] + nadh[c] + fum[c] <=> nad[c] + succ[c] ' |
| 39 | 'glucosamine 1 phosphate N acetyltransferase' | 'accoa[c] + gam1p[c] -> coa[c] + h[c] + acgam1p[c] ' |
| 40 | 'glycerol 3 phosphate dehydrogenase NAD ' | 'nad[c] + glyc3p[c] <=> h[c] + nadh[c] + dhap[c] ' |
| 41 | 'glycerol 3 phosphate dehydrogenase NADP ' | 'nadp[c] + glyc3p[c] <=> h[c] + nadph[c] + dhap[c] ' |
| 42 | 'UTP glucose 1 phosphate uridylyltransferase' | 'h[c] + utp[c] + g1p[c] <=> ppi[c] + udpg[c] ' |
| 43 | 'glutamine fructose 6 phosphate transaminase' | 'gln_L[c] + f6p[c] -> glu_L[c] + gam6p[c] ' |
| 44 | 'glutamine synthetase' | 'atp[c] + glu_L[c] + nh4[c] -> adp[c] + h[c] + pi[c] + gln_L[c] ' |
| 45 | '4 aminobutyrateglutamate antiport' | 'glu_L[e] + 4abut[c] <=> glu_L[c] + 4abut[e] ' |
| 46 | 'glutamate decarboxylase' | 'h[c] + glu_L[c] -> co2[c] + 4abut[c] ' |
| 47 | 'homocysteine S methyltransferase' | 'amet[c] + hcys_L[c] -> h[c] + ahcys[c] + met_L[c] ' |
| 48 | 'hypoxanthine phosphoribosyltransferase Hypoxanthine ' | 'prpp[c] + hxan[c] -> ppi[c] + imp[c] ' |
| 49 | 'isoleucine transaminase' | 'akg[c] + ile_L[c] <=> 3mop[c] + glu_L[c] ' |
| 50 | 'L isoeucine transport inout via proton symport' | 'h[e] + ile_L[e] <=> h[c] + ile_L[c] ' |
| 51 | 'inosine transport in via proton symport reversible' | 'h[e] + ins[e] <=> h[c] + ins[c] ' |
| 52 | 'L lactate dehydrogenase' | 'nad[c] + lac_L[c] + 1.125 pseud[c] <=> h[c] + nadh[c] + pyr[c] ' |
| 53 | 'L lactate reversible transport via proton symport' | 'h[e] + lac_L[e] <=> h[c] + lac_L[c] ' |
| 54 | 'methionine adenosyltransferase' | 'atp[c] + h2o[c] + met_L[c] -> pi[c] + ppi[c] + amet[c] ' |
| 55 | 'nucleoside diphosphate kinase ATPUDP ' | 'atp[c] + udp[c] <=> adp[c] + utp[c] ' |
| 56 | 'nucleoside diphosphate kinase ATPdCDP ' | 'atp[c] + dcdp[c] <=> adp[c] + dctp[c] ' |
| 57 | 'phosphoglucosamine mutase' | 'gam1p[c] <=> gam6p[c] ' |
| 58 | 'phosphoglucomutase' | 'g1p[c] <=> g6p[c] ' |
| 59 | 'inorganic diphosphatase' | 'h2o[c] + ppi[c] -> h[c] + 2 pi[c] ' |
| 60 | 'phosphopentomutase deoxyribose ' | '2dr1p[c] <=> 2dr5p[c] ' |
| 61 | 'phosphoribosylpyrophosphate synthetase' | 'atp[c] + r5p[c] <=> h[c] + amp[c] + prpp[c] ' |
| 62 | 'purine nucleoside phosphorylase Inosine ' | 'pi[c] + ins[c] <=> hxan[c] + r1p[c] ' |
| 63 | 'pyrimidine nucleoside phosphorylase uracil ' | 'pi[c] + uri[c] <=> ura[c] + r1p[c] ' |

| | | |
|---|---|---|
| 64 | 'ribokinase' | 'atp[c] + rib_D[c] -> adp[c] + h[c] + r5p[c] ' |
| 65 | 'ribosylhomocysteinase' | 'h2o[c] + rhcys[c] -> hcys_L[c] + rib_D[c] ' |
| 66 | 'ribonucleoside triphosphate reductase CTP ' | 'ctp[c] + trdrd[c] -> h2o[c] + dctp[c] + trdox[c] ' |
| 67 | 'L serine deaminase' | 'ser_L[c] -> pyr[c] + nh4[c] ' |
| 68 | 'L serine transport inout via proton symport' | 'h[e] + ser_L[e] <=> h[c] + ser_L[c] ' |
| 69 | 'succinate transporter inout via proton symport' | 'h[e] + succ[e] <=> h[c] + succ[c] ' |
| 70 | 'triose phosphate isomerase' | 'dhap[c] <=> g3p[c] ' |
| 71 | 'thioredoxin reductase NADPH ' | 'h[c] + nadph[c] + trdox[c] -> nadp[c] + trdrd[c] ' |
| 72 | 'UDP N acetylglucosamine diphosphorylase' | 'h[c] + utp[c] + acgam1p[c] -> uacgam[c] + ppi[c] ' |
| 73 | 'UDPglucose 6 dehydrogenase' | 'h2o[c] + 2 nad[c] + udpg[c] -> 3 h[c] + 2 nadh[c] + udpglcur[c] ' |
| 74 | 'uridylate kinase UMP ' | 'atp[c] + ump[c] -> adp[c] + udp[c] ' |
| 75 | 'uridine kinase ATPUridine ' | 'atp[c] + uri[c] -> adp[c] + h[c] + ump[c] ' |
| 76 | 'Ha out' | 'HA_monomer[e] -> ' |
| 77 | 'HA c2e' | 'HA_monomer[c] -> HA_monomer[e] ' |
| 78 | 'HAS' | 'uacgam[c] + udpglcur[c] -> 2 udp[c] + HA_monomer[c] ' |

**Figure S1:** Sub-network of over-expression targets from FSEOF analysis created using Escher (King et al., 2015)

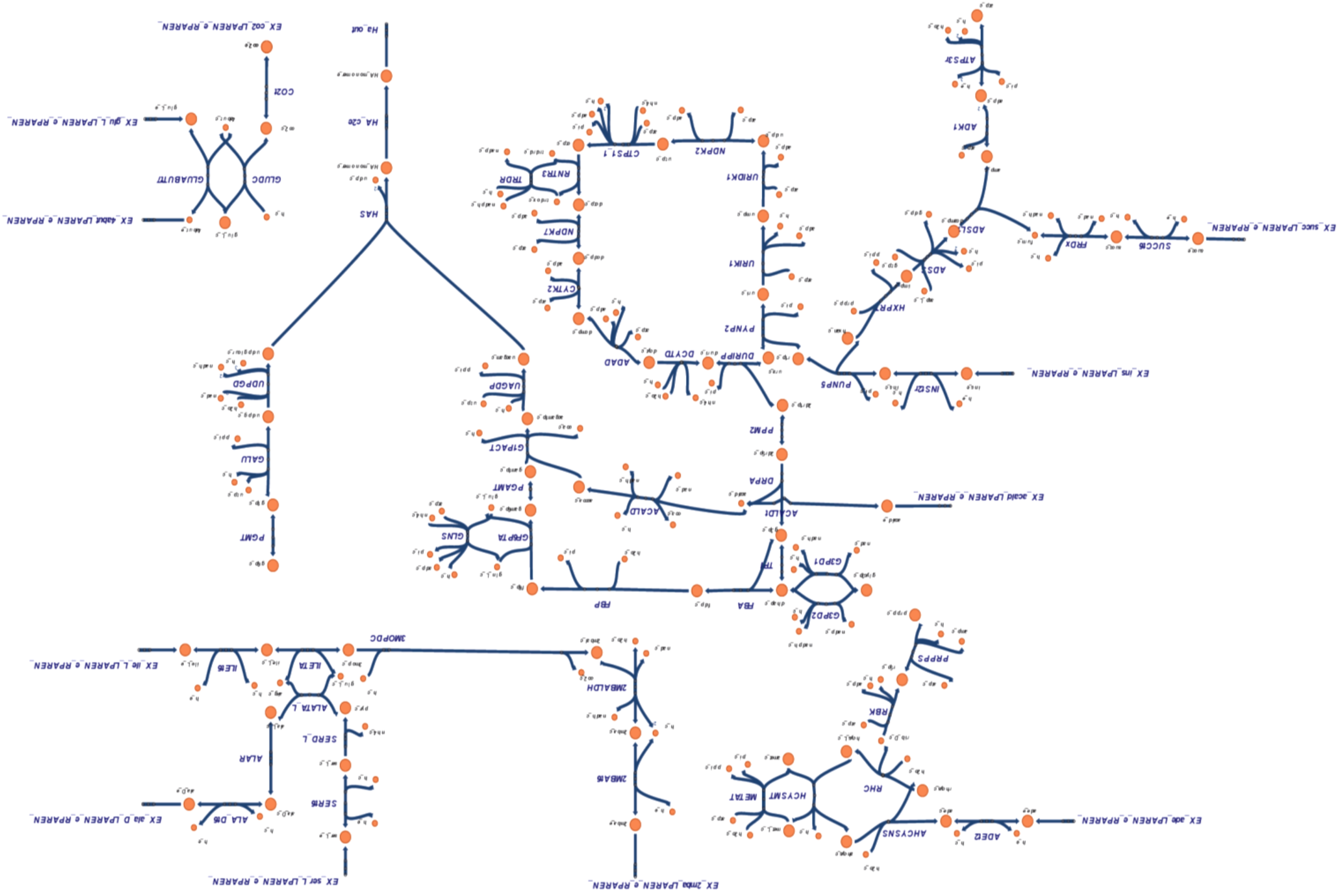

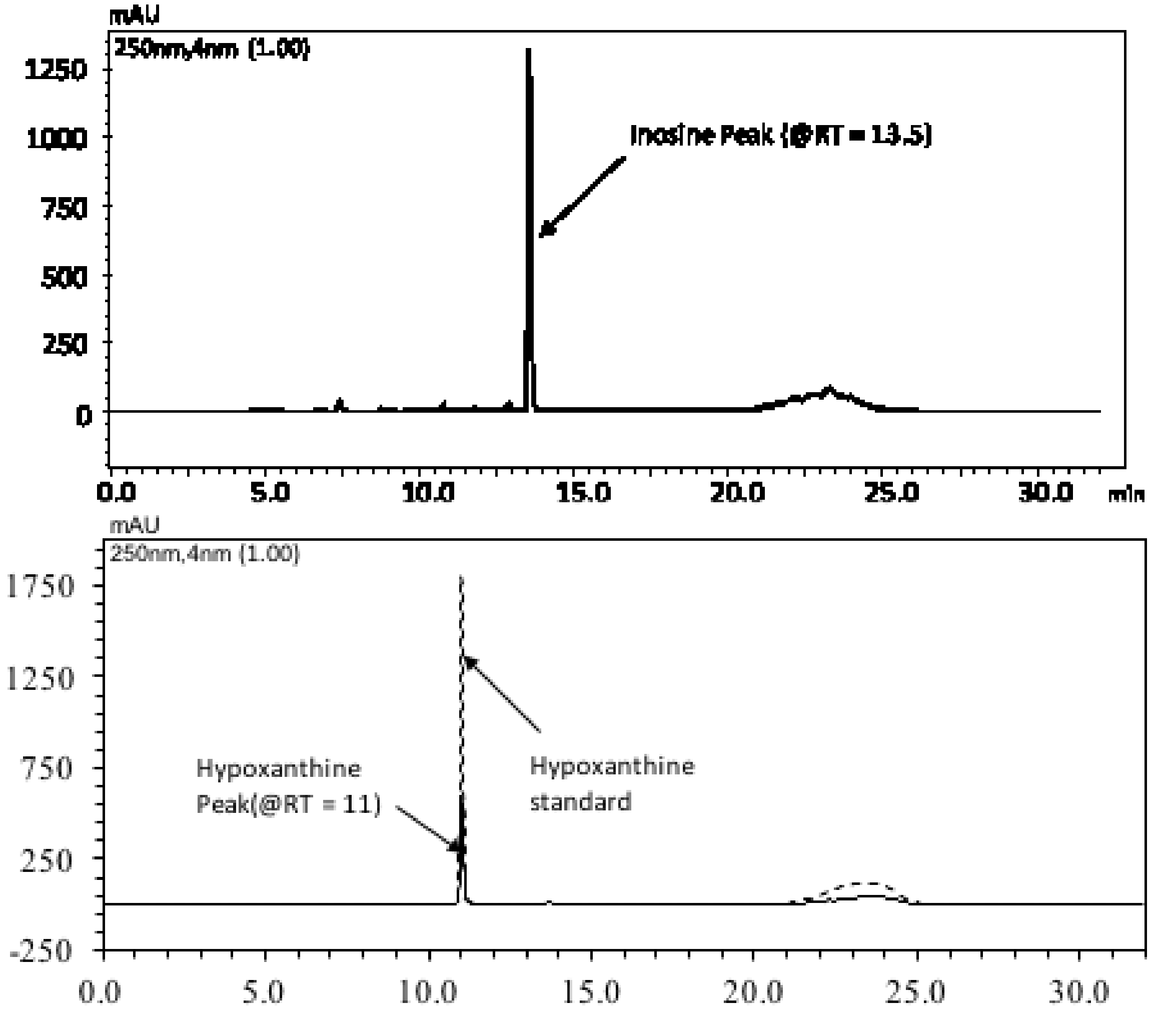


**Figure S2:** Reverse Phase Chromatogram of spent media at the (a) start with just inosine peak (RT = 13.5 mins) and (b) end with hypoxanthine (RT = 11 mins) and superimposed hypoxanthine standard (dashed) peaks

### A. Supplementary Methods

#### A.1 Standard plots and HPLC protocol

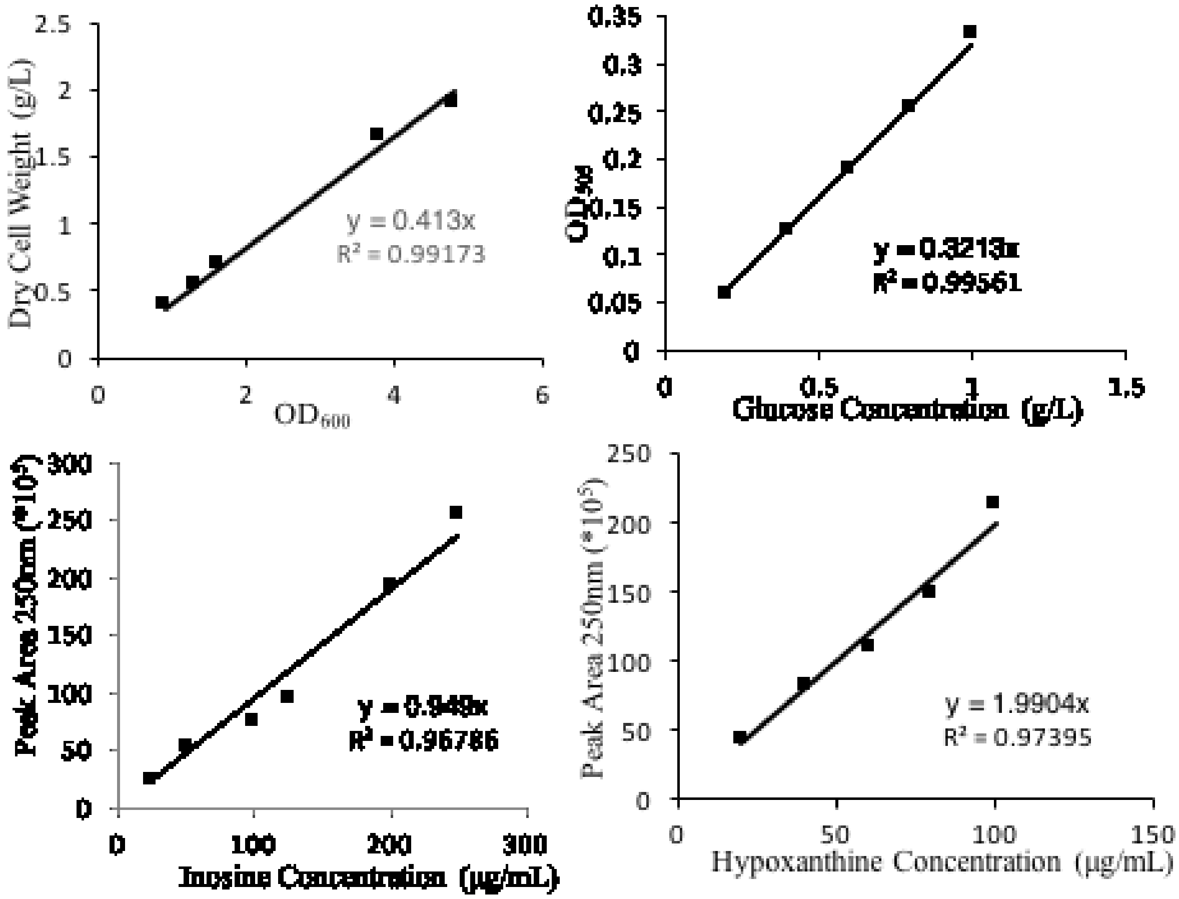


**Figure A.1.** Clockwise from top-left: Standard plots for Biomass, glucose, hypoxanthine and inosine estimation.

**Reverse Phase Gradient Protocol for the hypoxanthine and inosine estimation (adapted from (Farthing et al., 2007))**

- Monolithic Luna C-18 Phenomenex® column - length 250 mm, internal diameter of 4.6 mm, particle size of 5µ and pore size of 100 Å; Photodiode array (PDA) detector at 250 nm
- A flow rate of 0.6 mL/min
- Aqueous mobile phase - trifluoroacetic acid (0.05% TFA in deionized water pH 2.2, v/v); methanol gradient
- 32 minute time course per sample as follows: A - 0.05% TFA in deionized water; B - 100% Methanol -- 95:5::A:B (v/v) at 0 min ; 70:30::A:B (v/v) at 12 min; 10:90::A:B (v/v) at 13 min and held 3 min, and 95:5::A:B (v/v) at 17 min and hold for 15 minutes (to elute all other components)
- Inosine and Hypoxanthine elute at 13.5 and 11 mins respectively.

#### A.2 Adaptations to available *L. lactis* model

- Available *L. lactis* GSM (iNF518) 754 reactions, 650 metabolites and 518 genes.

- 35 exchange reactions with experimentally derived bounds were relaxed with default lower bounds of -1000 or 0 mmol/(g DCW·h) depending on reversibility and upper bounds of 1000 mmol/(g DCW·h).
- Hyaluronan synthase, HA transport and exchange reactions added -- 757 reactions, 652 metabolites and 519 genes (iNF519).

**Table A.1:** Reactions added to relaxed model to enable HA production

| Reaction Name | Reaction Formula |
|---|---|
| HA Synthase | UDP-glucuronic acid[c] + UDP-N-acetylglucosamine[c] → HA_Monomer[c] + 2UDP[c] |
| HA Transport | HA_Monomer[c] → HA_Monomer[e] |
| HA Exchange | HA_Monomer[e] ⇔ |

- **iNF519 + SJR6 chemostat data** for glucose consumption rate and lactate, acetate, ethanol and formate production rates (Badle et al., 2014)

**Table A.2:** Experimental flux bounds incorporated in the model

| Exchange Reaction | Lower bound (mmol/(g DCW·h)) | Upper bound (mmol/(g DCW·h)) |
|---|---|---|
| Glucose | -9.78 | -2.94 |
| Lactate | 3.44 | 14.1 |
| Acetate | 0.083 | 0.237 |
| Formate | 0 | 0.511 |
| Ethanol | 0.39 | 1.326 |

- **Model Cleaning**: iNF518 model -113 gaps in total (77 root gaps and 36 downstream gaps).

**Table A.3:** Reactions added and removed to the model

| S.No. | Reactions added |
|---|---|
| 1 | superoxide-forming NADH oxidase in *L. lactis* |
| 2 | Octaprenyl pyrophosphate synthase |
| 3 | Formation reaction for Acyl Carrier Protein (ACP) |
| 4 | Formation reaction for N-formylmethonine (fMet) |

| S.No. | Reactions Removed |
|---|---|
| 1 | undecaprenol kinase |
| 2 | N-hydroxyarylamine O-Acetyltransferase |
| 3 | 4-carboxymuconolactone decarboxylase |

| | |
|---|---|
| 4 | coproporphyrinogen oxidase 3 |
| 5 | hydroxybutyrate dehydrogenase |
| 6 | Arbutin 6-phosphate glucohydrolase |
| 7 | methylthioadenosine nucleosidase |
| 8 | myo-inositol-1-phosphatase |
| 9 | Glycerophosphodiester phosphodiesterase glycerophosphoinositol |
| 10 | Glycerophosphodiester phosphodiesterase glycerophosphoethanolamine |
| 11 | tetrahydropicolinate succinylase. |
| 12 | Amylomaltase maltotriose |
| 13 | Amylomaltase maltotetraose |
| 14 | Amylomaltase maltopentaose |
| 15 | Amylomaltase maltohexaose |
| 16 | CDP glycerol glycerophosphotransferase |
| 17 | CDP ribitol phosphoribitoltransferase |
| 18 | Chitinase |
| 19 | 2-dehydro-3-deoxy phosphogluconate aldolase |
| 20 | 5-formyltetrahydrofolate cyclo ligase |
| 21 | 5-methyltetrahydropteroyltriglutamate homocysteine S methyltransferase |
| 22 | Salicin-6-phosphate glucohydrolase. |

- Final model used in this study: 741 reactions, 618 metabolites, 519 genes and only 7 root gaps.

### B. Supplementary Results

#### B.1 Characteristics of in silico HA production

Following modifications of the model as described in §A.2, the model predicted *L. lactis* growth rate reasonably well (predicted growth rate of 0.322 $hr^{-1}$ for a chemostat run with steady state dilution rate of 0.3 $hr^{-1}$). It also proved to be capable of reproducing certain trends of HA flux that were previously reported in literature. For example, the model reproduced the positive correlation reported between HA and glucose (Badle et al., 2014; Prasad et al., 2010) for lower glucose uptake rates (Fig. B.1(a)). Simulations also showed an increase in HA with increase in glutamine uptake (Fig B.1(b)) as previously reported (Blank et al., 2005; Im et al., 2009; Shah et al., 2013). The simulations also indicated a decrease in HA flux with increase in lactate production flux or flux towards fructose- 6-phosphate (Fig. B.1(c) and (d) respectively) as reported earlier (Badle et al., 2014). Further, in silico HA flux increased with additional ATP formation flux (Fig. B.1(e)), thereby agreeing with previous reports on the energy-intensive characteristics of HA synthesis (Liu et al., 2008).

We also examined the theoretical maximum HA flux that can be produced under the given conditions of metabolite uptake, secretion and growth rates. Ideally, this is obtained by setting the product flux as the objective function. However, in this case, an objective function that maximized product yielded zero biomass, while that which maximized biomass yielded zero product flux. Due to this mutual exclusivity between biomass and HA fluxes, the conditions at which standard theoretical maximum flux is obtained is merely an impractical scenario.

To surmount this problem, the derivation of a theoretical maximum was slightly modified. To simulate a condition with a minimum non-zero growth, the maximum HA objective was augmented by fixing a lower bound for biomass corresponding to a lower growth rate of 0.1 $h^{-1}$ for which experimental values at steady state were available. With all other flux bounds retained at the same values, the maximum HA flux under these conditions was 19.107 mmol/(g DCW·h). This can be compared to the HA flux currently obtained in the recombinant *L. lactis* system at steady state in a CSTR with the 0.1 $h^{-1}$ dilution rate – 0.023 mmol/(g DCW·h). The order of magnitude difference in the theoretical maximum from the observed HA flux for this strain indicates that there is a huge room for improvement for the latter in this organism. The simulations also show a steady decrease in the theoretical maximum HA flux with increase in enforced minimum biomass flux (Fig. B.2). This indicates that a little compromise on the growth rate may drastically increase the theoretical maximum HA flux.

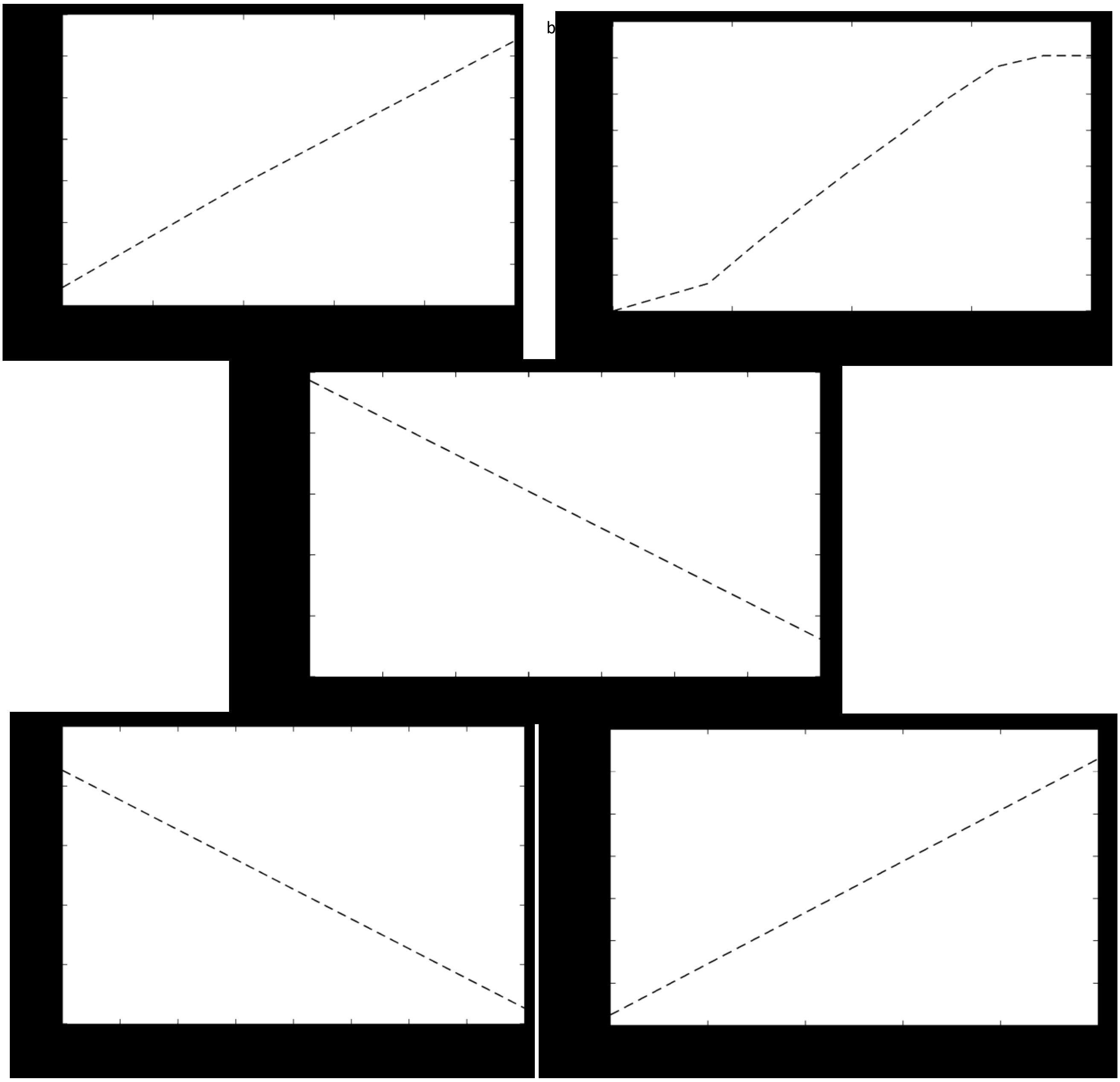


**Figure B.1:** Correlation between theoretical maximum HA flux and (a) glucose uptake (b) glutamine uptake (c) lactate production (d) fructose-6-phosphate production (e) additional ATP formation

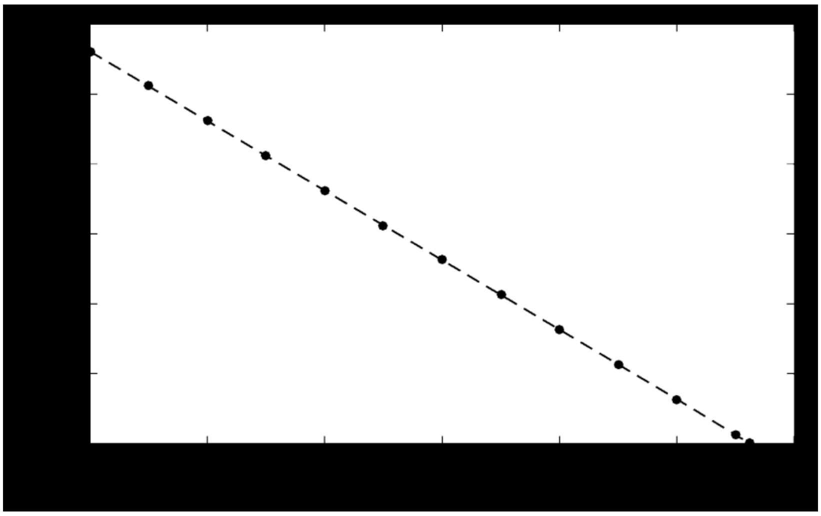

**Figure B.2:** Plot of predicted theoretical maximum HA flux for different biomass fluxes.

**B.2** ***Knock-outs predicted to increase the theoretical maximum***

Several algorithms have been developed until now for identification of knock-out targets (Burgard et al., 2003; Pharkya, 2004; Pharkya and Maranas, 2006; Tepper and Shlomi, 2010; Yang et al., 2011). The most widely used algorithm for this purpose is OptKnock, the bi-level platform that points out genes that need to be removed to simultaneously maximize growth rate and product flux (Burgard et al., 2003). Analyzing our model with OptKnock suggested three knock-out gene-targets, *viz.* lactate dehydrogenase (*ldh*), alcohol dehydrogenase (*adh*) and acetate kinase (*ak*) (Table B.1(a)). Out of these, only the *ldh* knock-out simulation showed a significant increase in the theoretical maximum HA and hence would be the foremost knock-out target derived out of this analysis. Previous reports on high titers of HA obtained from mutant strains that lack the *ldh* gene support this prediction (Kaur and Jayaraman, 2016).

**B.3** ***Knock-outs to reach the theoretical maximum***

This section stems from the order-of-magnitude difference between the theoretical maximum HA flux and those obtained at steady state in the organism. The difference indicates that knock-outs that take the system closer to the theoretical maximum from the current state have the potential to make a bigger impact on HA flux than those predicted to increase theoretical maximum HA flux (in §B.2). To identify these, we listed out a set of reactions that are switched off (fluxes go to zero from a non-zero value) when shifted from conditions maximizing biomass to those maximizing HA. The major reactions that were put forth as knock-out targets in this category belonged to Glycolysis and Pentose Phosphate Pathway (Table B.1(b)). This signifies that the major roadblock in achieving theoretical maximum HA is the competition for carbon from pathways that are related to growth and energy production.

**Table B.1: (a)** Knock-outs to increase theoretical maximum (OptKnock) **(b)** Knock-outs to reach theoretical maximum

(a)

| S. No. | Enzyme Knocked out | ΔTheoretical maximum |
|---|---|---|
| 1. | Lactate dehydrogenase | 0.167 |
| 2. | Alcohol dehydrogenase | 0.004 |
| 3. | Acetate Kinase | E^-5 |

(b)

| Knock-outs to reach theoretical maximum |
|---|
| Glucose-6-phosphate dehydrogenase |
| Phosphogluconate dehydrogenase |
| 6-phosphogluconolactonase |
| Phosphofructokinase |
| L-threonine transport |
